# Compatibility of time-lapse dry incubator on *in vitro* production of bovine embryos

**DOI:** 10.1101/2024.08.01.606272

**Authors:** Haruhisa Tsuji, Hiroki Nagai, Sayaka Kobinata, Hinata Koyama, Atchalalt Khurchabilig, Noritaka Fukunaga, Yoshimasa Asada, Satoshi Sugimura

**Author notes:** **Corresponding author:** (S. Sugimura).

## Abstract

Embryo culture is crucial to achieve successful outcomes in assisted reproductive technology (ART) for cattle. This study explored the innovative use of dry incubators integrated with time-lapse monitoring systems for bovine embryo culture, building on their advantages in human medicine, such as reduced contamination risk, stable temperature control, and lower gas consumption. Our research demonstrates the feasibility of this approach, showing that although the osmotic pressure gradually increases over the culture period, it remains below the critical threshold for developmental impairment. Embryos cultured in dry incubators exhibited morphokinetics comparable to those cultured in conventional humidified time-lapse incubators. Furthermore, RNA-seq revealed that the transcriptomic profiles of blastocysts cultured in dry incubators closely matched those of blastocysts cultured in humidified incubators. These findings highlight the significant potential of dry incubators with time-lapse monitoring systems for the in vitro production of bovine embryos, marking a promising advancement in assisted reproductive technologies for the livestock industry and research setting.

## 1. Introduction

Assisted reproductive technology (ART) has garnered worldwide attention due to its potential for facilitating genetic gains in livestock breeding. Within this landscape, *in vitro* production (IVP) has emerged as a prominent technique offering precise control of reproductive processes. This is particularly evident in bovine reproduction, in which IVP is a crucial tool for advancing reproductive research and improving livestock breeding programs. Parallel to advancements in human medicine, there has been a notable shift towards the use of IVP techniques in bovine reproduction. This transition is fueled by the desire to enhance breeding efficiency, overcome infertility challenges, and accelerate genetic progress in cattle populations. As a result, the demand for innovative technologies that can support the culture of bovine embryos with optimal developmental outcomes is growing [1-3].

In human ART, the integration of dry incubators with time-lapse imaging systems is gaining traction as a high-quality assessment platform. Dry incubators offer distinct advantages over traditional humidified incubators, including a reduced risk of contamination, stable temperature control, and minimal maintenance requirements, including gas consumption (carbon dioxide and nitrogen) [4-6]. These features make dry incubators an attractive option for embryologists and clinicians, particularly for high-throughput ART laboratories.

Time-lapse imaging enables the continuous monitoring of embryo development, facilitating the assessment of morphokinetic parameters and dynamic changes in embryo behavior. This approach has shown promise in improving embryo selection based on developmental dynamics and morphological features, ultimately enhancing clinical outcomes [7-9]. Recently, automatic analysis has been introduced for embryo assessment, further improving the accuracy and efficiency of embryo selection by leveraging advanced algorithms and machine learning techniques [10].

Generally, the assessment of bovine embryo quality is performed by morphological grading on days 7 to 8, as recommended by the International Embryo Technology Society (IETS) [11,12]. However, the pregnancy success rate of embryos judged as transferable (Code 1 or 2) is only 30–50%, and conventional assessment lacks objectivity due to differing evaluations between operators [13,14]. To address this issue, time-lapse monitoring has emerged as a valuable tool for evaluating embryo quality during bovine ART. By tracking the developmental progression of bovine embryos in real-time, operators can assess developmental milestones, cell division patterns, and embryo kinetics, providing insights into embryo viability and selection criteria [14-18].

Despite these advancements, the compatibility of dry incubators with time-lapse monitoring systems for bovine embryo cultures remains uncertain. Therefore, in this study, we explored the feasibility of using dry incubators for bovine embryo culture and their compatibility with time-lapse imaging systems. To address this knowledge gap, we aimed to elucidate whether dry incubators could support both the culture and dynamic assessment of bovine embryos, thereby advancing our understanding of embryo development and quality assessment in agricultural and research settings.

## 2. Materials and Methods

### 2.1. Chemicals

All chemicals were purchased from Sigma-Aldrich (St. Louis, MO, USA) unless otherwise specified.

### 2.2. Osmotic pressure of culture media

The osmolality of the culture medium was measured using an osmometer (OSMOMATO 3000D; Gonotec, Berlin, Germany). The equipment was calibrated with 0, 300, and 800 mOsm kg^−1^ solutions with sample sizes of 50 µL in 500-µL measuring tubes.

### 2.3. Oocyte collection

Ovaries from Japanese Black or Japanese Black × Holstein breeds were collected from a local slaughterhouse and transported to the laboratory of the Tokyo University of Agriculture and Technology, Fuchu Campus, in an insulated container within 2 h. After transportation, the ovaries were washed and stored in physiological saline solution containing 0.9% NaCl at 38.0°C. Cumulus oocyte complexes (COCs) were aspirated from small follicles (2–6 mm in diameter) using a 10-mL syringe equipped with a 19-gauge needle.

### 2.4. *In vitro* maturation

The *in vitro* maturation (IVM) of bovine COCs was performed as previously described [15]. The IVM medium consisted of 25 mM 4-(2-hydroxyethyl) piperazine-1-ethanesulfonic acid-buffered tissue culture medium 199 (M199; Gibco, Paisley, Scotland, UK) supplemented with 5% calf serum (CS; Gibco) and 0.1 IU/mL recombinant human follicle-stimulating hormone (GONAL-F; Merck, Darmstadt, Germany). The COCs were washed with IVM medium and then incubated in 100 μL of IVM medium (20–25/droplet) covered with sterile paraffin oil (Nakarai Tesque, Kyoto, Japan) in 60-mm culture dishes (Nunc; Nalge Nunc International, Roskilde, Denmark). COCs were placed in a CO_2_ incubator (Astec, Fukuoka, Japan) at 38.5°C for 20 h in a humidified atmosphere with 5% CO_2_.

### 2.5. *In vitro* fertilization

*In vitro* fertilization was performed as previously described [15]. After 20 h of IVM, frozen sperm obtained from Japanese Black bulls was thawed and centrifuged in 90% Percoll solution at 740 × *g* for 10 min. After centrifugation, the pellet was resuspended in sperm washing medium, which consisted of Brackett and Oliphant (BO) solution [19] supplemented with 20 mM hypotaurine and 10 μL/mL heparin (Novo-Heparin Injection 1000; Aventis Pharma Ltd., Tokyo, Japan). The suspension was centrifuged at 540 × *g* for 5 min. The pellet was then resuspended in sperm washing medium and BO solution supplemented with 20 mg/mL bovine serum, and the final concentration was adjusted to 3 × 10^6^ spermatozoa/mL. Droplets of this suspension (100 μL each) placed in 60-mm dishes and covered with paraffin oil served as fertilization droplets. COCs were placed in fertilization droplets (20 COCs/droplet) and cultured for 6 h at 38.5°C in a humidified atmosphere with 5% CO_2_ (Astec).

### 2.6. *In vitro* culture of zygotes

Zygotes were washed several times with BO-IVC medium (IVF Biosciences). BO-IVC medium (100 μL) was placed within the circular wall of microwell culture dish (LinKID micro25; Dai Nippon Printing Co., Ltd., Kashiwa, Japan) and covered with 4.5 mL of paraffin oil (Nakarai Tesque) [15]. Twenty-five zygotes were placed in the microwells of a microwell culture dish (one zygote per microwell). *In vitro* culture was performed at 38.5°C in 5% O_2_, 5% CO_2_, and 90% N_2_ for 168 h in a humidified incubator (CCM-MULTI, Astec / APM-30D; Astec) and a dry incubator (CCM-iBIS; Astec).

### 2.7. Morphological quality of blastocysts

Blastocyst diameter was measured using a stereomicroscope equipped with a 5.0 μg/ml Hoechst 33342 and observed using a fluorescent microscope (BZ-X810; KEYENCE, Osaka, Japan). In addition, the embryos were classified into Code 1 to 3 according to the percentage of degenerated cells based on the criteria recommended by international embryo technology society (IETS) [11,12]: (i) Code 1: irregularities should be relatively minor and at least 85% of the cellular material should be an intact, viable embryonic mass; (ii) code 2: at least 50% of the cellular material should be an intact, viable embryonic mass; (iii) code 3: at least 25% of the cellular material should be an intact, viable embryonic mass.

### 2.8. Time-lapse monitoring

Morphokinetics were analyzed using a humidified incubator with time-lapse cinematography (CCM-MULTI; Astec) or a time-lapse dry incubator (CCM-iBIS; Astec) (Fig. 1 and Supplementary Movie 1). During the 168-h culture period, photographs of the zygotes were taken at 15-min intervals using an objective with 10× magnification. Morphokinetic parameters were annotated, including the timing of blastomere cleavage (t2 to t8), the start of the morula stage (tM) and blastulation (tSB), the formation of full blastocyst (tB), the expansion of blastocysts (tEB), and the start of hatching (tHB) after IVC was started.

**Fig. 1.**
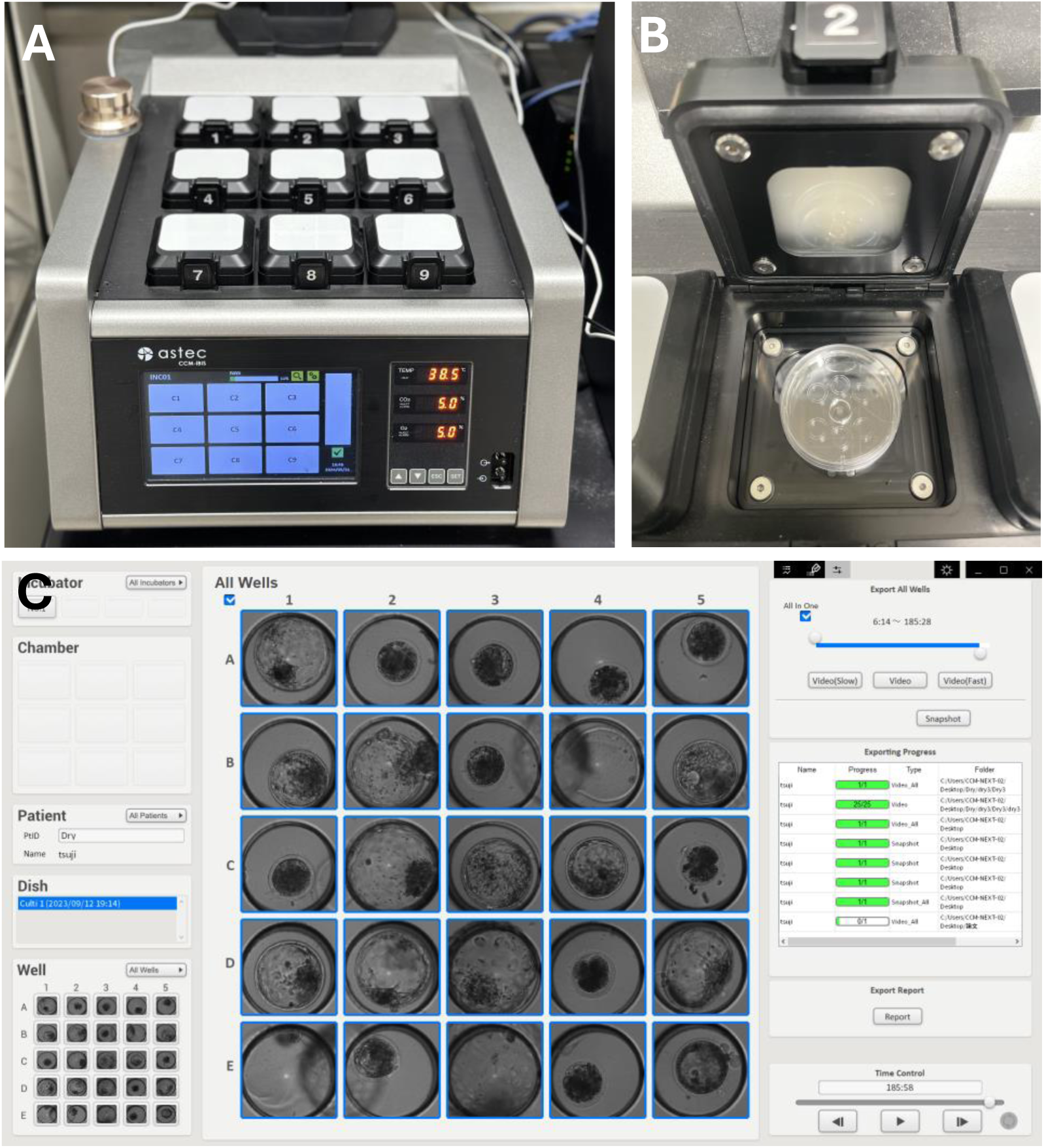
Photographs of the time-lapse dry incubator (CCM-iBIS; Astec). It has nine individual chambers, each of which allows for time-lapse observation. The external dimension of the main unit is 382 × 590 × 219 mm (width × diameter × height) (**A**). A specially developed well of the well (WOW)-type culture dish (LinKID^Ⓡ^ micro25; Dai Nippon Printing Co., Ltd.) [15] is placed in each chamber (**B**). Screen shot of LinKID^Ⓡ^ image viewer (Dai Nippon Printing Co., Ltd.) on day 8 of culture. The morphokinetics in each microwell can be observed in real time (see Suppl. Movie 1) (**C**).

### 2.9. RNA-seq

The zona pellucida was removed from all blastocysts using 0.25% pronase (Actinase E; Kakenseiyaku, Japan) to prevent contamination with nucleic acids from cumulus cells and sperm. Total RNA extraction, reverse transcription, and cDNA library construction were performed using a SMART-Seq HT PLUS Kit (Takara Bio). The quality of cDNA was assessed using an Agilent 2200 TapeStation (Agilent Technologies, Santa Clara, CA, USA) with a High Sensitivity D5000 ScreenTape (Agilent Technologies). Sequencing was performed on a NovaSeq 6000 platform (Illumina, San Diego, CA, USA) using 150 bp × 2 paired-end reads. Adapter trimming was performed using Trim-galore, quality checks were conducted using fastqc, mapping was performed using STAR with ARS-UCD1.2/bosTau9 as the reference, and expression quantification was performed using RSEM. Expression data (total counts) were then imported into iDEP.96 and were first normalized (CPM, count per million), while data transformation was conducted using EdgeR (log2(CPM+c)) as described previously [20]. mRNAs with a minimum expression of 0.5 CPM (count per million) expression in at least one sample were retained. Hierarchical clustering and principal component analysis (PCA) were performed using the correlation distance and average linkage. DESeq2 was used to identify Differential Expressed Genes (DEGs). The FDR cutoff was 0.1 and the minimum fold change was 2. The DEGs were visualized using Manhattan plots.

### 2.9. Statistical analysis

All data were analyzed using R. *P* < 0.05 was considered statistically significant. All percentage data were arcsine transformed.

## 3. Results

### 3.1. Osmotic instability of culture medium when using a dry incubator

Studies using human culture media have reported that while humidified incubators can maintain osmotic pressure, they are unstable in dry incubators [21]. Similarly, in this study, the humidified incubator maintained stable osmolarity levels around 270 mOsm from the beginning (263.0 ± 1.0 mOsm) until 8 days of culture (271.7 ± 2.1 mOsm). In contrast, the osmotic pressure in the dry incubator gradually increased, reaching 299 ± 15.4 mOsm on day 8 of culture. At all time points, except on day 8, osmolality was significantly higher under dry conditions than under humidified conditions.

### 3.2. Comparison of developmental competence and morphological quality of bovine embryos cultured in humid and dry incubators

No significant differences were observed in the cleavage and blastocyst formation rates between the humidified and dry incubators (Table 1). Additionally, there was no significant difference in morphological quality including the mean ± SD of diameter (237.4 ± 82.6 and 242.8 ± 73.3, respectively), the mean ± SD of total number of cells (189.2 ± 82.6 and 242.8 ± 73.3, respectively), and the percentage of IETS grading (Code 1: 40.3 and 42.0; Code 2: 50.0 and 42.0; Code 3: 9.7 and 15.9, respectively) between the humidified and dry incubators (Fig. 2).

**Fig. 2.**
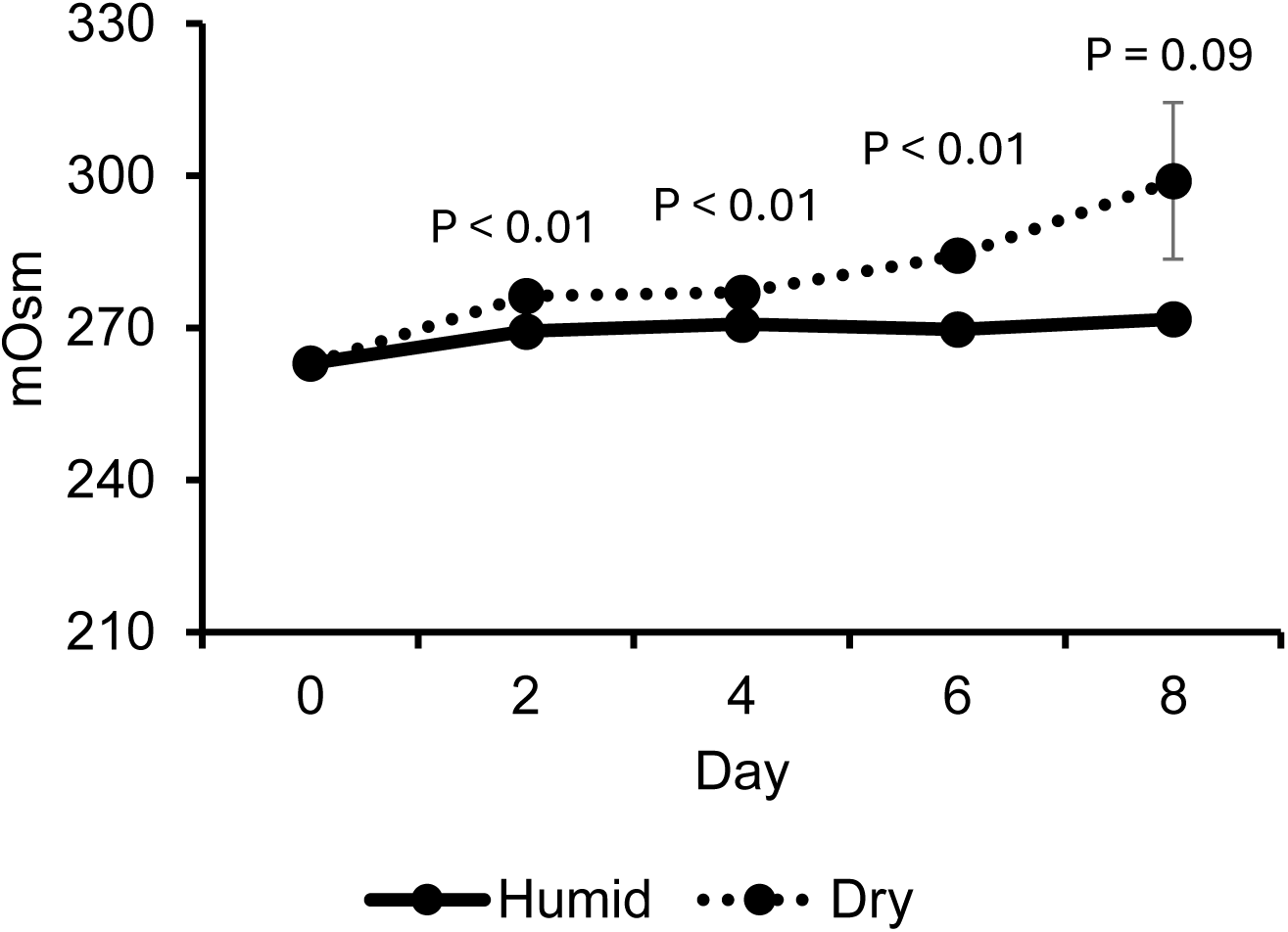
Temporal changes in the osmotic pressure of the IVC medium cultured in a humidity and dry incubators. Values represent the mean ± SD of three replicates.

**Table 1.** Effect of the dry incubator on in vitro development of bovine zygotes ^a^.

| Incubator<br>type | No. of<br>culture<br>zygotes | No. of embryos (%) |  |  |
| --- | --- | --- | --- | --- |
|  |  | Cleaved | Developed to blastocyst stage |  |
|  |  |  | Day 7 | Day 8 |
| Humid | 125 | 104 (85.0±6.8) | 63 (60.1±3.8) | 65 (62.4±1.2) |
| Dry | 125 | 107 (86.0±5.2) | 65 (61.9±5.5) | 73 (69.8±7.7) |

<sup>a</sup> Values represent the mean ± SD of four replicates.

### 3.4. Culturing in a dry incubator had no negative impact on the morphokinetics of bovine embryos

Dry conditions have been reported to delay the development of human embryos, but their effects on cattle remain unknown [4]. No significant differences were observed in the morphokinetic parameters of zygotes cultured in humid and dry incubators, except for tM. Interestingly, tM was faster in the dry incubator for approximately 5 h (Table 2).

**Table 2.** Timing of developmental event of embryos cultured in humidity and dry incubators.

|  | Humid incubator |  |  | Dry incubator |  |  | *p*-value |
| --- | --- | --- | --- | --- | --- | --- | --- |
|  | n | Mean | SD | n | Mean | SD |  |
| t2 | 62 | 17.56 | 3.50 | 69 | 18.54 | 2.19 | 0.062 |
| t4 | 62 | 26.04 | 3.57 | 69 | 27.09 | 3.22 | 0.081 |
| t6 | 62 | 33.69 | 3.42 | 69 | 34.72 | 3.89 | 0.114 |
| t8 | 62 | 35.85 | 2.58 | 69 | 36.91 | 3.64 | 0.054 |
| t9 | 62 | 65.76 | 15.45 | 69 | 68.26 | 12.87 | 0.318 |
| tM | 62 | 93.14 | 9.20 | 69 | 88.55 | 9.13 | 0.005 |
| tSB | 62 | 128.60 | 11.35 | 69 | 132.07 | 11.57 | 0.087 |
| tB | 62 | 140.34 | 11.58 | 69 | 142.51 | 13.44 | 0.328 |
| tEB | 41 | 152.11 | 7.38 | 48 | 154.78 | 9.59 | 0.141 |
| tHB | 12 | 155.71 | 4.37 | 19 | 159.42 | 7.11 | 0.119 |

t2–t8, time of cleavage to 2–9 cells; tM, time of the formation of fully compact morula; tSB, time of the start of blastulation; tB, time of the formation of full blastocyst; tEB, time of the formation of expanded blastocyst; tHB, time of the start of hatching. All timings are expressed as hours after IVC starting.

### 3.3. Transcriptomic profile in blastocysts cultured dry incubator

No clustering was observed between the humid and dry incubators in the hierarchical clustering and PCA (Fig. 4A and B). DEG analysis showed that 18 genes were upregulated and five genes were downregulated in the dry incubator (Fig. 4C). Interestingly, two of the upregulated genes, *TKDP1* [22,23] and *SSLP1* [24] were related to the differentiation and proliferation of the trophectoderm, and both were significantly more highly expressed in the dry incubator (Fig. 4D). No enrichment based on the DEGs gene set was observed in the GO or KEGG analyses. These results suggested similarities in the transcriptional profiles of bovine blastocysts cultured in humidified and dry incubators.

**Fig. 3.**
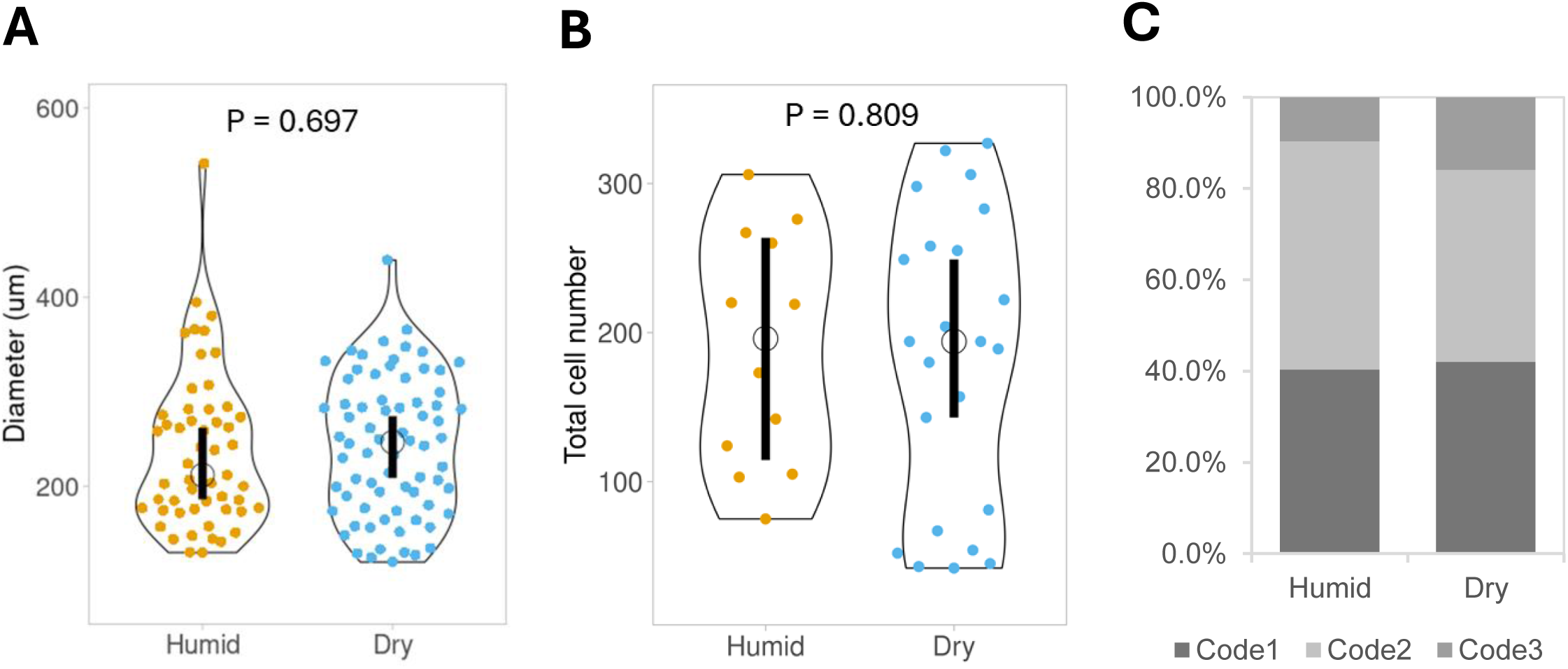
Morphological quality of blastocysts at day 8 cultured in humidity and dry incubators. The diameter (**A**) and total cell number (**B**) are represented by a violin plot, with medians shown as circles and 95% CI as black bars. Population of Code 1, 2, and 3 (IETS criteria): (i) Code 1: irregularities should be relatively minor and at least 85% of the cellular material should be an intact, viable embryonic mass; (ii) Code 2: at least 50% of the cellular material should be an intact, viable embryonic mass; (iii) Code 3: at least 25% of the cellular material should be an intact, viable embryonic mass.

**Fig. 4.**
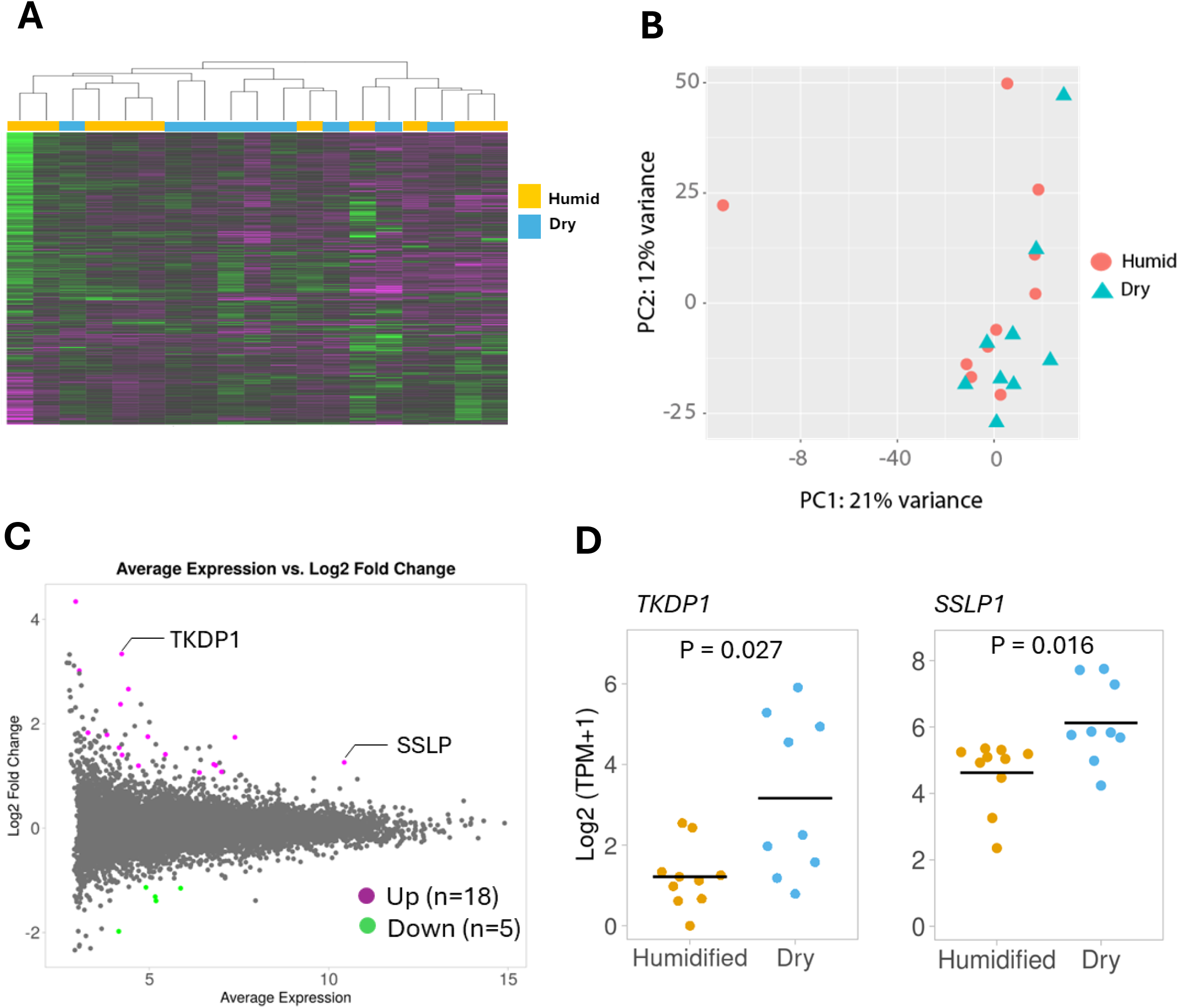
RNA-seq analysis of blastocysts at day 8 cultured in humidity and dry incubators. Heatmap with hierarchical clustering analysis showing top 1000 most variable genes (**A**) and two-dimensional principal component analysis (PCA) (**B**) from RNA-seq data. The x-axis of the Manhattan plot represents average expression and y-axis represents log_2_ fold change value. The significant up- and down-differentially expressed genes (DEGs) in blastocysts cultured in dry incubator are presented in violet and green, respectively (**C**). The expression of genes (TKDP1 and SSLP1) related to differentiation and proliferation of trophectoderm identified as DEGs. Black bars indicate median values (**D**).

## 4. Discussion

In this study, we investigated the feasibility of utilizing dry incubators for bovine embryo culture and their compatibility with time-lapse monitoring systems, with the aim of contributing to the advancement of assisted reproductive technologies (ART) in bovine reproduction. Our findings shed light on several key aspects of the use of dry incubators.

One notable finding from our study is the stability of the osmotic pressure in the culture medium when using dry incubators compared to humidified incubators. While humidified incubators maintained stable osmolarity levels throughout the culture period, osmotic pressure gradually increased in dry incubators, resulting in significantly higher levels by the end of the culture period. Previous reports have indicated that >300 mOsm adversely affects embryonic development [25-27] and epigenetics [28]. Our results showed that the osmotic pressure gradually increased over 8 d of dry incubator cultivation, reaching nearly 300 mOsm by day 8. However, both incubation conditions resulted in similar cleavage and blastocyst formation rates and no significant differences in blastocyst morphological quality. These observations suggest that despite the gradual increase in osmotic pressure in the dry incubator, the pressure remained within a range that did not significantly affect bovine embryo development. It is possible that the bovine embryos were able to tolerate slight fluctuations in osmotic pressure without compromising their developmental competence.

However, these findings emphasize the importance of close monitoring and regulation of osmotic pressure during embryo culture, particularly when using dry incubators [21,29]. It has been reported that increasing the amount of oil [29] or using heavy oil can reduce the increase in osmolarity [21]. Using a medium with a low osmolarity is also effective in reducing the negative effects of osmolarity [29]. CR1aa with 5% calf serum, which is widely used in bovine embryo production, had a high osmolarity of 273 mOsm (data not shown), whereas the BO-IVC used in this study had a relatively low osmolarity of 263 mOsm. In dry incubators, selecting a medium with low osmolarity, such as BO-IVC, may facilitate better outcomes. Our results demonstrate that osmotic pressure does not exceed the reported threshold associated with developmental impairment; thus, maintaining stable osmotic conditions remains crucial for optimizing embryo culture conditions and ensuring consistent developmental outcomes.

This study highlights an interesting observation in bovine embryos, in which the overall morphokinetics were similar between embryos cultured in humidified and dry incubators. Contrary to the results observed in bovine embryos, previous reports on humans have indicated that developmental kinetics are delayed when cultured in a dry environment [4]. This discrepancy suggests species-specific differences in responses to environmental conditions, such as osmotic stress.

In addition, RNA-seq analysis revealed that the overall transcriptional profiles of bovine blastocysts cultured in dry and humidified incubators were remarkably similar. Interestingly, this study found an elevated expression of genes related to trophectoderm differentiation and proliferation, such as *TKDP1* [22,23] and *SSLP1* [24], in blastocysts cultured in a dry incubator [22]. This specific upregulation aligns with previous reports showing that osmotic stress promotes trophectoderm development [31]. A dry incubator likely induces mild osmotic stress, which can act as a physiological signal to enhance the differentiation and proliferation of the trophectoderm, contributing to enhanced conceptus elongation. This further supports the suitability of dry incubators for bovine embryo cultures.

Overall, our study provides valuable insights into the use of dry incubators for bovine embryo culture and their compatibility with time-lapse imaging. However, further research is needed to fully elucidate the long-term effects after transfer and the optimal conditions for utilizing dry incubators for cattle reproduction.

## 5. Conclusion

In this study, dry incubators with time-lapse imaging systems were demonstrated to represent a viable option for the culture and selection of bovine embryos, maintaining developmental outcomes similar to those of humidified incubators. These findings support the feasibility of the use of time-lapse dry incubators for advancing bovine-assisted reproductive technologies in agricultural and research settings.

## Acknoledgments

This work was supported by JSPS KAKENHI Grant Number JP23K23760 to S.S., the JRA Livestock Industry Promotion Project to S.S., and TUAT’s Competitive Research Program TAMAGO to S.S. We would like to thank Editage (www.editage.com) for English language editing.

## Author contributions

**Haruhisa Tsuji:** Conceptualization, Methodology, Investigation, Data curation, Formal analysis, Writing – original draft, Visualization. **Hiroki Nagai:** Methodology. Ayaka Kobinata: Methodology. Hinata Koyama: Methodology. **Atchalalt Khurchabilig:** Methodology. **Noritaka Fukunaga:** Writing – review & editing. **Yoshimasa Asada:** Writing – review & editing. **Satoshi Sugimura:** Conceptualization, Methodology, Writing – original draft, Writing – review & editing, Supervision, Funding acquisition, Project administration.

