## Supplementary movie legend for "Compatibility of time-lapse dry incubator on *in vitro* production of bovine embryos"

**Supplementary Information**

**Supplementary Movie legends**

**Supplementary Movie 1.** Time-lapse monitoring of bovine IVP zygotes. This movie corresponds to Figure 1C. Images were captured at 15-min intervals for eight days.
